# “The genome assembly of the duckweed fern, *Azolla caroliniana*”

**DOI:** 10.1101/2024.10.22.619683

**Authors:** Michael J. Song, Fay-Wei Li, Forrest Freund, Merly Escalona, Erin Toffelmier, Courtney Miller, H. Bradley Shaffer, Oanh Nguyen, Mohan P. A. Marimuthu, Noravit Chumchim, Carrie Tribble, Colin W. Fairbairn, William Seligmann, Carl J. Rothfels

## Abstract

*Azolla* is a genus of freshwater ferns that is economically important as a nitrogen-fixing biofertilizer, biofuel, bioremediator, and for potential carbon sequestration, but also contains weedy invasive species. In California, only two species are currently recognized but there may be up to six putative species, with the discrepancy being due to the difficulty in identifying taxa, hybridization, and the introduction of non-native species. Here, we report a new haplotype-resolved, chromosome-level assembly of *Azolla caroliniana* as part of the California Conservation Genomics Project (CCGP), using a combination of PacBio HiFi and Omni-C sequencing technologies. The assembly is 521 Mb in length, with a contig N50 of 1.6 Mb, and is scaffolded into 22 pseudo-chromosomes. The BUSCO completeness score is 87.5%, making it the most complete and most contiguous *Azolla* assembly to date. In combination with the previously published *A. filiculoides* genome, this *A. caroliniana* genome will be a powerful tool for understanding the population genetics and taxonomy of one of the most cryptic, economically important, and poorly circumscribed fern taxa, and for facilitating land plant genomics more broadly.

## Introduction

*Azolla* Lam. is a genus of tiny, heterosporous, floating freshwater ferns with an outsized importance (Figure 1). These ferns harbor obligate, endosymbiotic, nitrogen-fixing cyanobacteria in leaf pockets (Li et al., 2018, de Vries and de Vries, 2022) and have been used for centuries as a green biofertilizer, often in rice production (Lumpkin and Plucknett, 1982). *Azolla* has been touted as having great economic and ecological potential as a biofertilizer, phytoremediator, and biofuel (Brouwer et al. 2014), and also as a method of carbon sequestration to combat climate change—the *Azolla* Event during the Eocene may have sequestered enough carbon to transform the hot-house planet of the Paleocene–Eocene Thermal Maximum to the ice-house planet that we have today (Speelman et al., 2009). However, some species of *Azolla* are federally listed noxious weeds (USDA Federal Noxious Weed List) and complicating this matter is the lack of consensus on how many species there are, how to identify them, and what to name them. Additionally, species in this genus are known to hybridize, and there are many known instances of polyploidy (Watanabe et al., 1993; Stergianou and Fowler, 1990).

**Figure 1.**
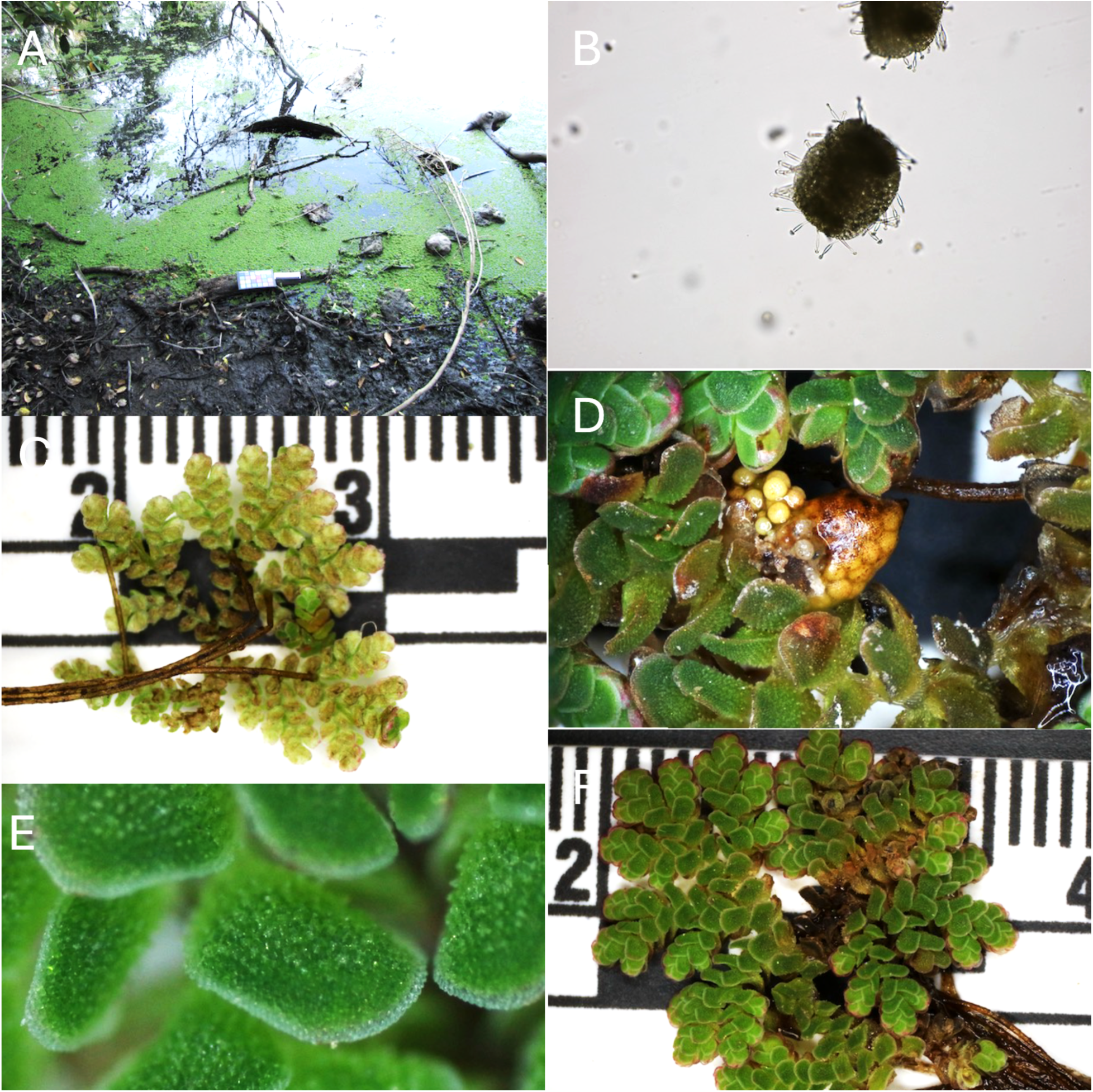
*Azolla caroliniana* from the genome-sample collection site: Lake Anza, Tilden Regional Parks, Contra Costa County, California. (A) Habitat; (B) Microsporocarp with glochidia; (C) Whole-plant abaxial surface; (D) Megasporocarps showing megaspores; (E) Close up of adaxial leaf surface; (F) Whole-plant adaxial surface. Photo credit: Forrest Freund.

In California, while there are only two currently recognized species (Jepson Flora Project 2024), up to five described species occur (*A. caroliniana, A. filiculoides, A. mexicana, A. microphylla,* and *A. rubra*), plus least one additional undescribed species (Rothfels and Li, unpublished), and the recently introduced *A. pinnata* ssp. *pinnata* (Song et al. 2023). The taxonomy of the group is both complicated and controversial (Edvard and Van Hove, 2004; Madeira et al. 2013). By developing genomic resources for *Azolla*, we can begin to understand which species are present, how they can be distinguished, what their distributions and abundances are, as well as identify centers of genomic diversity and patterns of connectivity. These genomic resources will also help investigate broader evolutionary questions such as the origins of heterospory, diversification of the microbiome, and evolution of symbioses and organelles.

Here we report a haplotype-resolved, chromosome-level genome assembly for *Azolla caroliniana* Willd., produced as part of the California Conservation Genomics Project (CCGP). The goal of the CCGP is to assemble and annotate high quality genomes of 150 species across the state in conjunction with whole-genome resequencing data to identify diversity hotspots and inform conservation and management plans (Shaffer et al., 2022; Toffelmier et al., 2022; Fiedler et al. 2022; Beninde et al. 2022). The *A. caroliniana* genome will be a powerful tool for understanding the landscape genomics and taxonomy of one of the most cryptic, economically important, noxious, and poorly circumscribed fern taxa.

## Methods

### Biological Materials

#### Field collections and identification

A collection that we originally identified as *A. microphylla* (Tilden East Bay Regional Park; 37.89685, -122.24980, Freund 365) was chosen to serve as the reference genome for *Azolla* in the CCGP. However, analysis of the genome resequencing data revealed that many of our samples (including this reference) are nested in groups usually treated as *A. caroliniana* (Song et al., 2024) and therefore we treat this sample as *A. caroliniana* in this paper. The voucher specimen is located at the University Herbarium (Herbarium code: UC) at the University of California Berkeley.

#### Surface sterilization

We followed a modified version of the surface-sterilization protocol in Dijkhuizen (2018) as described in Song et al. (2024).

### Genome Assembly

#### DNA extraction, library preparation, and sequencing

We extracted high molecular weight genomic DNA (gDNA) from 158 mg of whole plant tissues using the cetyltrimethylammonium bromide (CTAB) method as described in Inglis et al. (2018), with the following modifications: (i) we used sodium metabisulfite (1% w/v) instead of 2-mercaptoethanol in the sorbitol wash buffer and CTAB solutions; (ii) we repeated the tissue homogenate wash steps until the supernatant turned clear; (iii) we performed the CTAB lysis step at 45°C and; (iv) performed the chloroform extraction step twice using ice-cold chloroform. The DNA yield (650 ng) was quantified using a Quantus Fluorometer (QuantiFluor ONE dsDNA Dye assay; Promega, WI), and the size distribution of the DNA was estimated using the Femto Pulse system (Genomic DNA 165 kb kit, Agilent, CA), where 70% of the DNA fragments were found to be 30 kb or longer.

The HiFi SMRTbell library was constructed using the SMRTbell gDNA Sample Amplification Kit (Pacific Biosciences [PacBio], Menlo Park, CA; Cat. #101-980-000) and the SMRTbell Express Template Prep Kit 2.0 (PacBio Cat. #100-938-900) according to the manufacturer’s instructions. Approximately 10 kb sheared DNA by the Megaruptor 3 system (Diagenode, Belgium; Cat. #B06010003) was used for removal of single-strand overhangs at 37°C for 15 minutes, DNA damage repair at 37°C for 30 minutes, end repair and A-tailing at 20°C for 30 minutes and 65°C for 30 minutes, and ligation of overhang adapters at 20°C for 60 minutes. To prepare for library amplification by PCR, the library was purified with ProNex beads (Promega, Madison, WI; Cat. # NG2002) for two PCR amplification conditions at 15 cycles each then another ProNex beads purification. Purified amplified DNA from both reactions were pooled in equal mass quantities for another round of enzymatic steps that included DNA repair, end repair/A-tailing, overhang adapter ligation, and purification with ProNex Beads. The PippinHT system (Sage Science, Beverly, MA; Cat # HPE7510) was used for SMRTbell library size selection to remove fragments < 6–10 kb. The 10–11 kb average HiFi SMRTbell library was sequenced at UC Davis DNA Technologies Core (Davis, CA) using one 8M SMRT cell, Sequel II sequencing chemistry 2.0, and 30-hour movies, on a PacBio Sequel IIe sequencer.

The Omni-C library was prepared using the Dovetail Omni-C Kit (Dovetail Genomics, Scotts Valley, CA) according to the manufacturer’s protocol with slight modifications. First, specimen tissue (whole plant) was thoroughly ground with a mortar and pestle while cooled with liquid nitrogen. Nuclear isolation was then performed using the method of Workman et al. (2018). Subsequently, chromatin was fixed in place in the nucleus and digested under DNase I until a suitable DNA fragment-length distribution was obtained. Chromatin ends were repaired and ligated to a biotinylated bridge adapter followed by proximity ligation of adapter-containing ends. After proximity ligation, crosslinks were reversed and the DNA was purified from proteins. Purified DNA was treated to remove biotin that was not internal to ligated fragments. An NGS library was generated using an NEB Ultra II DNA Library Prep kit (NEB, Ipswich, MA) with an Illumina compatible y-adaptor. Biotin-containing fragments were then captured using streptavidin beads. The post-capture product was split into two replicates prior to PCR enrichment to preserve library complexity with each replicate receiving unique dual indices. The library was sequenced at Vincent J. Coates Genomics Sequencing Lab (Berkeley, CA) on an Illumina NovaSeq 6000 platform (Illumina, CA) to generate approximately 100 million 2 ✕ 150 bp read pairs per GB genome size.

All data generated by CCGP can be found in the NCBI SRA (PRJNA720569).

#### Nuclear genome assembly

We assembled the genome of the *Azolla caroliniana* sample following the CCGP assembly pipeline Version 5.0, as outlined in Table 1. We removed the remnant adapter sequences from the PacBio HiFi dataset using HiFiAdapterFilt (Sim et al. 2022) and generated an initial diploid phased assembly using HiFiasm (Cheng et al. 2021, Cheng et al, 2022) in HiC mode with the filtered PacBio HiFi reads and the OmniC short reads, a process that generates two assemblies, one per haplotype. We then aligned the Omni-C data to both assemblies following the Arima Genomics Mapping Pipeline (https://github.com/ArimaGenomics/mapping_pipeline) and scaffolded both assemblies with SALSA (Ghurye et al. 2017, 2019).

**Table 1:**
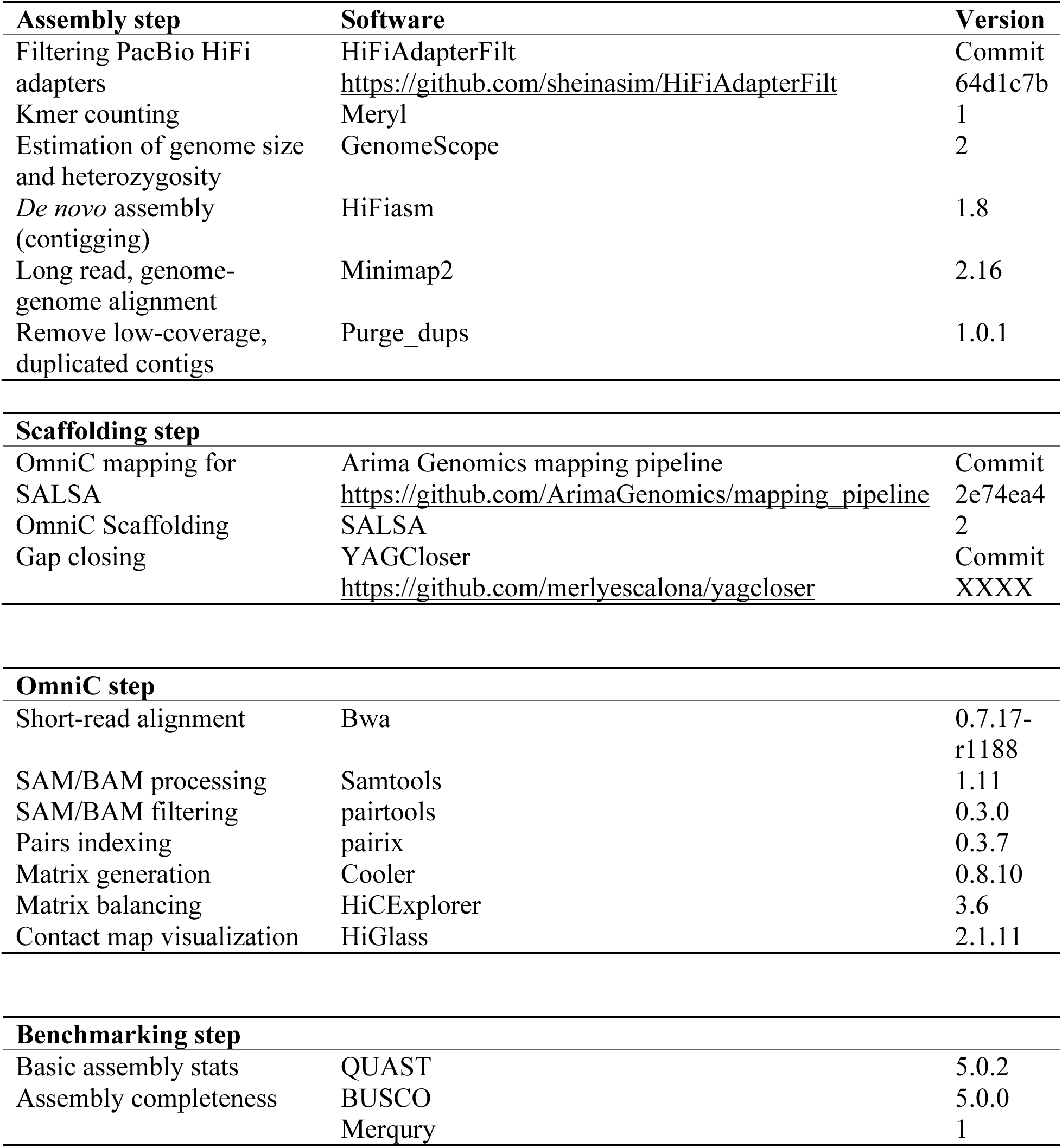
Assembly pipeline and software used. Software citations are found in the references.

The assemblies were manually curated by iteratively generating and analyzing their corresponding Omni-C contact maps. In general, to generate the contact maps we aligned the Omni-C data with BWA-MEM (Li 2013), identified ligation junctions and generated Omni-C pairs using pairtools (Lee et al. 2022; Open2C et al. 2023). Then, we generated multi-resolution Omni-C matrices with cooler (Abdennur and Mirny 2020) and balanced them with hicExplorer (Ramírez et al. 2018). We used HiGlass (Kerpedjiev et al. 2018) and the PretextSuite (https://github.com/wtsi-hpag/PretextView; https://github.com/wtsi-hpag/PretextMap; https://github.com/wtsi-hpag/PretextSnapshot) to visualize the contact maps where we identified misassemblies and misjoins, and finally modified the assemblies using the Rapid Curation pipeline from the Wellcome Trust Sanger Institute, Genome Reference Informatics Team (https://gitlab.com/wtsi-grit/rapid-curation). Some of the remaining gaps (joins generated during scaffolding and/or curation) were closed using the PacBio HiFi reads and YAGCloser (https://github.com/merlyescalona/yagcloser). We checked for contamination using the BlobToolKit Framework (Challis et al. 2020) and tagged final assemblies as primary and alternate haplotype assemblies, following overall quality metrics, where the primary haplotype assembly is the assembly that is more complete.

#### Genome quality assessment

We generated k-mer counts from the PacBio HiFi reads using meryl (https://github.com/marbl/meryl). The k-mer counts were then used in GenomeScope2.0 (Ranallo-Benavidez et al. 2020) to estimate genome features including genome size, heterozygosity, and repeat content. To obtain general contiguity metrics, we ran QUAST (Gurevich et al. 2013). To evaluate genome quality and functional completeness we used BUSCO (Manni et al. 2021) with the Viridiplantae ortholog database (viridiplantae_odb10), which contains 425 genes. Assessment of base level accuracy (QV) and k-mer completeness was performed using the previously generated meryl database and merqury (Rhie et al. 2020). We further estimated genome assembly accuracy via BUSCO gene set frameshift analysis using the pipeline described in Korlach et al. (2017). Measurements of the size of the phased blocks is based on the size of the contigs generated by HiFiasm on HiC mode. We follow the quality metric nomenclature established by Rhie et al. (2021), with the genome quality code x.y.P.Q.C, where, x = log10[contig NG50]; y = log10[scaffold NG50]; P = log10 [phased block NG50]; Q = Phred base accuracy QV (quality value); C = % genome represented by the first “n” scaffolds, following a karyotype of 2n=44 for this species, as found in the Genome on a Tree database (Accessed Sept. 3rd, 2024; Challis et al. 2023). Quality metrics for the notation were calculated on the assembly for the haplotype that is more complete. Dotplot comparing *A. caroliniana* and *A. filiculoides* genomes (Li et al., 2018) was done by D-Genies (Cabanettes and Klopp, 2018).

#### Chloroplast Genome

We assembled the chloroplast genome of *A. caroliniana* from the PacBio HiFi reads using Oatk (https://github.com/c-zhou/Oatk). We used GeSeq (Tillich et al. 2017) to generate a draft genome annotation and searched for matches of the resulting chloroplast assembly sequence in the nuclear genome assembly for this sample using BLAST+ (Camacho et al. 2009) and filtered out contigs based on matches with a breadth of coverage of the organelle of 100% and a percentage of sequence identity >99%. The assembly was manually curated by joining the sequences corresponding to the typical chloroplast genome structure.

#### Sample identification and phylogenetic analyses

In order to confirm that our sample was *A. caroliniana* and not the expected *A. microphylla,* we performed several phylogenetic analyses in addition to the analyses of Song et al. (2024), which used SNP data and plastome trees. For these analyses we used a set of previously sampled *Azolla* species from Li et al. (2018; *Azolla nilotica* SRR6482158; *Azolla caroliniana* SRR6480231 and SRR6480201; *Azolla microphylla* SRR6480161; *Azolla mexicana* SRR6480159; and *Azolla filiculoides* SRR6932851) and 22 CCGP resequencing samples (Freund 403, Freund 408, Freund 410, Freund 411, Freund 425, Freund 435, Freund 444, Freund 445, Freund 446, Freund 448,

Freund 454, Freund 458, Freund 469, Freund 477, Freund 485, Freund 487, Freund 488, Freund 490, Freund 491, Freund 494, Freund 502, Freund 507, Freund 508). The CCGP samples were selected to be representative of the major clades found in California (Song et al., 2024). For each sample, we used the CAPTUS phylogenomics pipeline (Ortiz et al., 2023) with default parameters to perform de novo assembly, extract the default Angiosperms353 markers (Johnson et al., 2019), and perform multi-species alignment (“mafft_auto”; Katoh et al., 2005).. We inferred a maximum likelihood phylogeny from these concatenated data with IQTree v2.3.5 (Minh et al., 2020), using the partition model -p, where each gene partition is allowed to have its own average substitution rate (Chernomor et al., 2016). We also used IQTree to infer individual gene trees for each locus using the -S option. These 330 gene trees were used as input for species-tree inference using ASTRAL v5.7.8, which accounts for incomplete lineage sorting (Zhang et al., 2018).

## Results

### Sequencing data

The Omni-C library generated 81.34 million read pairs and the PacBio HiFi library generated 3.3 million reads. The PacBio HiFi sequences yielded ∼54-fold genome coverage and had an N50 read length of 9,262 bp; a minimum read length of 109 bp; a mean read length of 9,167 bp; and a maximum read length of 34,900 bp (see Supplementary Fig. 1 for read length distribution). Based on the PacBio HiFi data, Genomescope 2.0 estimated a genome size of 556.06 Mb, a 0.18 % sequencing error rate, 1.19% heterozygosity and k-mer uniqueness of 45.7%. The k-mer spectrum shows a bimodal distribution with a major coverage peak at ∼50-fold coverage and a minor coverage peak ∼26 coverage (Figure 2A).

**Figure 2.**
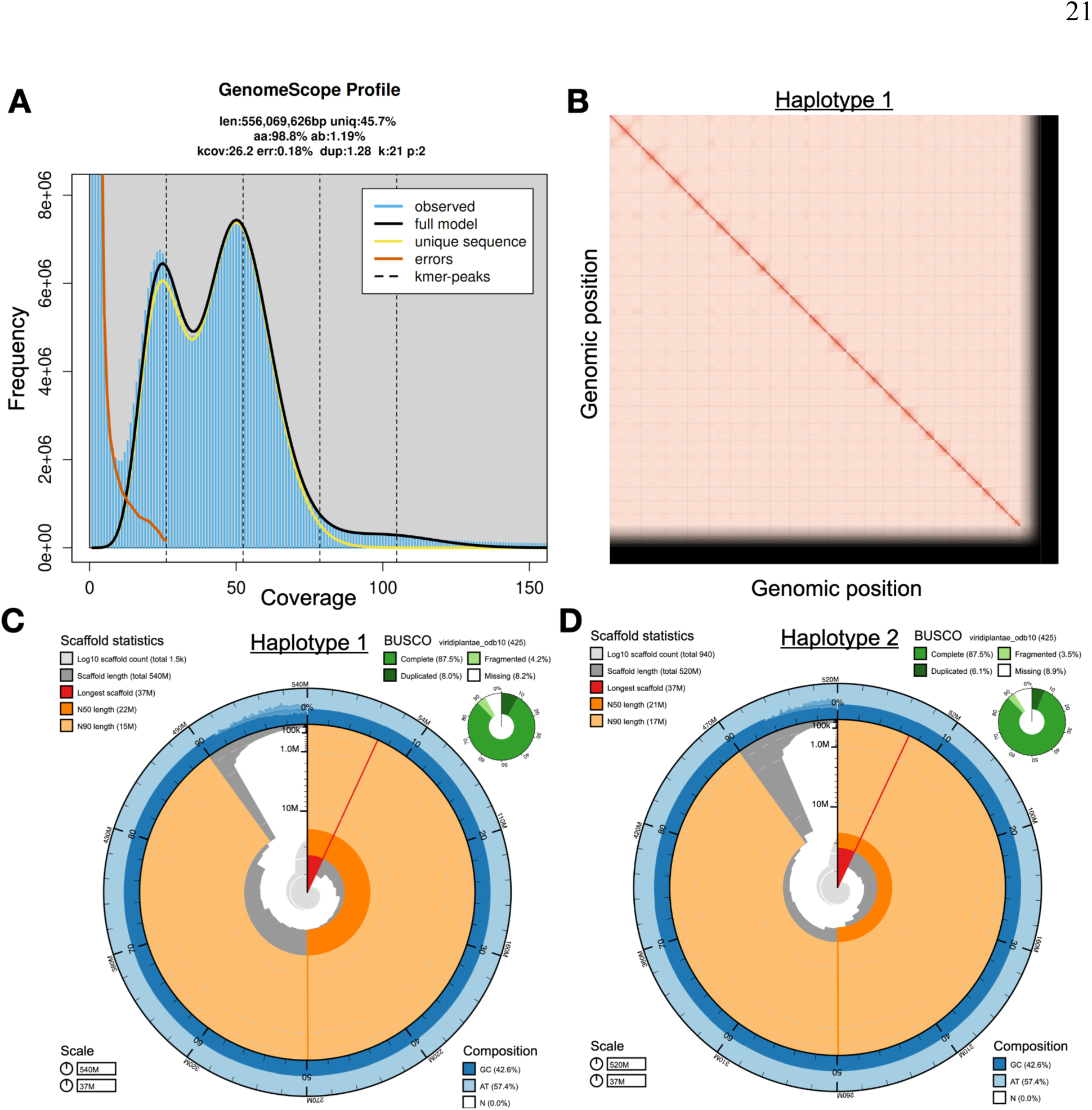
*Azolla caroliniana* chromosome-level genome assembly. (A) K-mer spectra output generated from PacBio HiFi data without adapters using GenomeScope2.0. The bimodal pattern observed corresponds to a diploid genome and the k-mer profile matches that of high heterozygosity. K-mers covered at lower coverage and high frequency correspond to differences between haplotypes, whereas the higher coverage and slightly lower frequency correspond to the similarities between haplotypes. (B) Omni-C Contact maps for the curated genome assembly of haplotype 1 generated with PretextSnapshot. Omni-C contact maps translate proximity of genomic regions in 3D space to contiguous linear organization. Each cell in the contact map corresponds to sequencing data supporting the linkage (or join) between two of such regions. Scaffolds are separated by black lines. (C) Haplotype 1 and (D) haplotype 2 BlobToolKit snail plot showing a graphical representation of the quality metrics presented in Table 2. The first diagonal (∼30 degrees) line represents the size of the longest scaffold; all other scaffolds are arranged in order of size clockwise around the plot and drawn in gray starting from the outside of the central plot. Dark and light orange arcs show the scaffold N50 and scaffold N90 values. The central light gray spiral shows the cumulative scaffold count with a white line at each order of magnitude. White regions in this area reflect the proportion of Ns in the assembly; the dark vs. light blue area around it shows mean, maximum and minimum GC vs. AT content at 0.1% intervals (Challis et al. 2020).

**Table 2:**
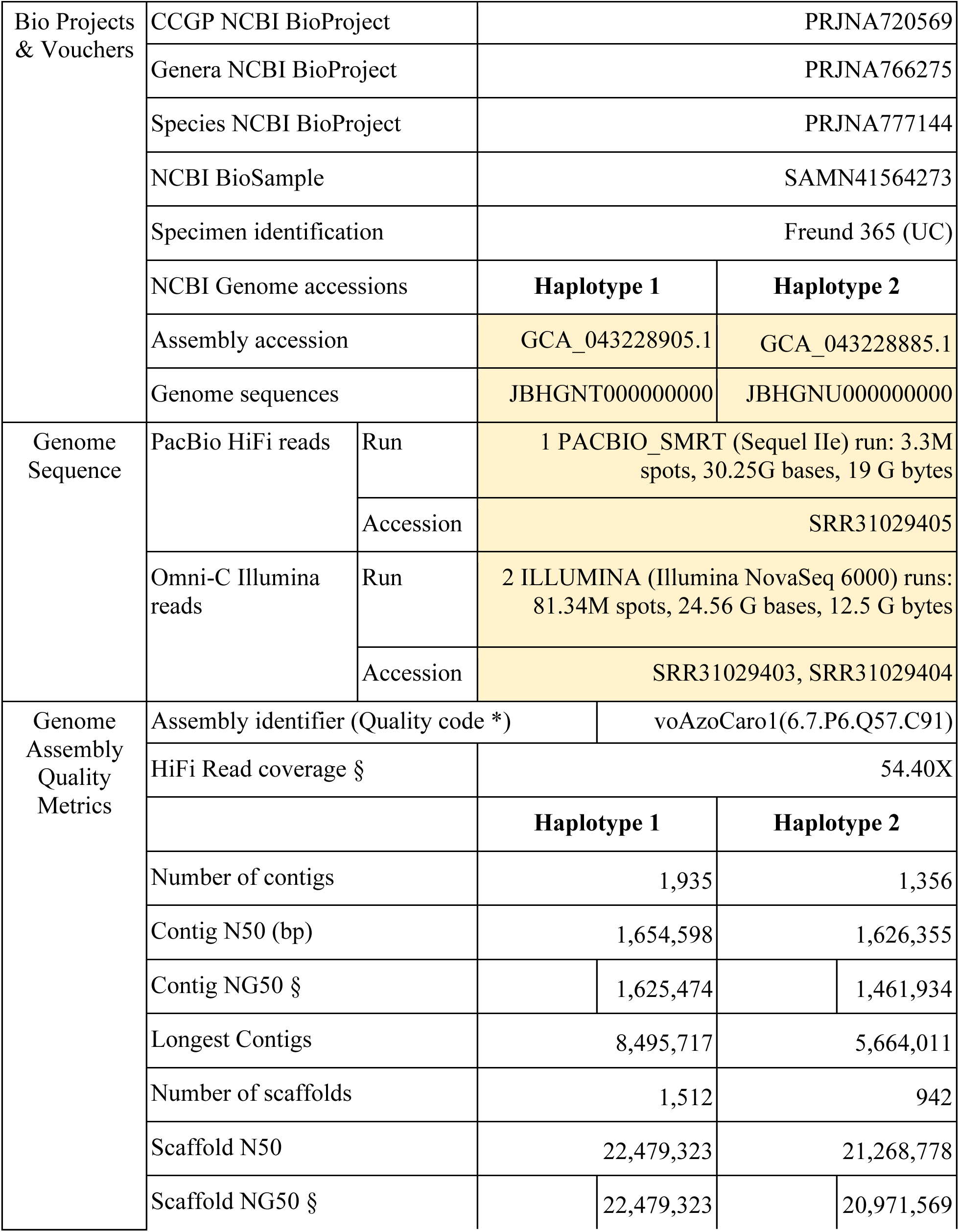
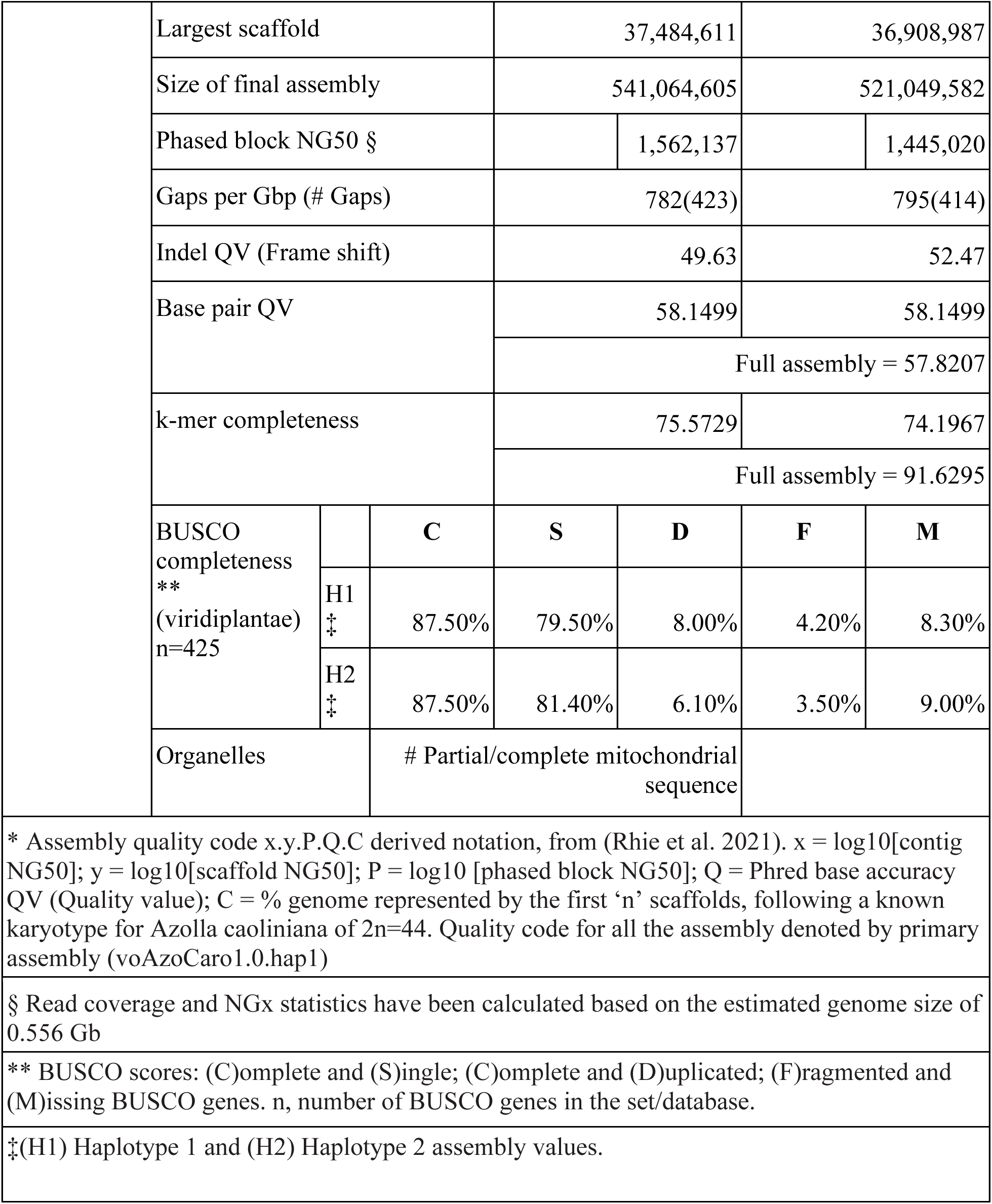
Sequencing and assembly statistics, and accession numbers.

### Nuclear genome assembly

The final genome assembly (voAzoCaro1) consists of two phased haplotypes. Both assemblies are similar in size, with a difference of ∼20 Mb between haplotypes. The final haplotype assemblies are also similar in size to the estimated genome assembly size from GenomeScope2.0.

The primary assembly (voAzoCaro1.0.p) consists of 1,503 scaffolds spanning 523.24 Mb with a contig N50 of 1.68 Mp, a scaffold N50 of 22.24 Mb, the largest contig size of 8.49 Mb, and the largest scaffold size of 37.34 Mb. The alternate assembly (voAzoCaro1.0.a) consists of 942 scaffolds spanning 504.25 Mb with a contig N50 of 1.62 Mb, a scaffold N50 of 21.26 Mb, the largest contig size of 5.66 Mb, and the largest scaffold size of 36.25 Mb.

The primary assembly has a BUSCO completeness score for the Viridiplantae gene set of 84.7%, a base pair quality value (QV) of 56.98, a kmer completeness of 73.39%, and a frameshift indel QV of 49.63. The alternate assembly has a BUSCO completeness score for the same gene set of 83.1%, a base pair QV of 57.45, a kmer completeness of 71.66%, and a frameshift indel QV of 52.47.

During manual curation we made a total of 155 joins (94 on the primary assembly and 61 on the alternate assembly) and 60 breaks (39 on the primary assembly and 21 on the alternate assembly) based on the signal from the Omni-C contact maps. We filtered out a total of 353 contigs based on the contamination screening with Blobtools. After inspection upon submission to NCBI, we removed nine contigs (detailed taxonomic assignment of removed contigs can be found on Supplementary Table 1–3). We also filtered out nine contigs corresponding to chloroplast contamination (one from the primary assembly and eight from the alternate assembly). No other contigs were removed.

The Omni-C contact maps for both assemblies show highly contiguous assemblies (Figure 2B). Detailed assembly statistics are reported in Table 2 and represented graphically in Figure 2C. We have deposited the genome assembly on NCBI GenBank (See Table 2 and Data Availability for details).

*Azolla* species are reported to harbor not only the obligate endosymbiont *Anabaena azollae*, but also a suite of other bacterial endophytes in the leaf cavities (Li et al., 2018; Dijkhuizen et al., 2018; Song et al. 2024). Indeed, a large number of contigs from the initial HiFi assembly were classified as bacterial, whose GC content and/or coverage deviated considerably from the plant contigs (Supplementary Fig. 1). Compared to the previously published contig-level assembly of *A. filiculoides*, this *A. caroliniana* genome is considerably more heterozygous and slightly smaller in size. On the other hand, the two genomes are largely collinear (Supplementary Fig. 2).

### Chloroplast genome assembly

The final (partial) chloroplast sequence has a size of 149,725 bp, where the LSC spans 83,717 bp, the SSC 24,426 bp and the IRs 20,776 bp. Regions are separated by 10bp gaps. Final annotation consists of 14 rRNAs, 51 transfer RNAs, and 96 protein coding genes. Our partial assembly was 149 kb, close to the 147 kb reported for *A. filiculoides* (Accession: MF177094.1), and is thus likely to be nearly complete"

### Taxon identification

Our reference genome comes out nested within *Azolla caroliniana* in both the concatenated-data tree and the ASTRAL species tree and therefore we treat this taxon as *Azolla caroliniana* in this paper (Supplementary. Fig. 3).

## Discussion

We provide the most complete *Azolla* genome to date and the second chromosome-level assembly in Salviniales (Rahmatpour et al., 2023). Our *A. caroliniana* genome assembly was scaffolded onto 22 pseudochromosomes, which is the expected haploid number of *Azolla* (Fig. 2B; Flora of North America). While being largely collinear, the *A. caroliniana* assembly is slightly smaller than the *A. filiculoides* assembly (540/521 Mb vs 623 Mb). This difference might be due to variation in repeat content and/or the way contamination was filtered. As shown in Supplementary Fig. 1, *Azolla* contains a diversity of endophytes, the genomes of which were assembled alongside the host genome. The different methods and thresholds used to distinguish plant vs non-plant contigs could lead to a size difference, especially considering that Hi-C contact information was not incorporated in assembling the *A. filiculoides* genome.

One striking difference between the two *Azolla* genomes is the level of heterozygosity—high in *A. caroliniana* and low in *A. filiculoides*. The strain used for *A. filiculoides* sequencing has gone through a few rounds of inbreeding for the purpose of turning it into a lab strain (Brouwer et al., 2014), hence the low heterozygosity. On the other hand, the *A. caroliniana* sample was collected straight from the field. It is worth noting that both haplotypes of *A. caroliniana* have high contig N50 and were scaffolded onto 22 pseudochromosomes, thus representing the first diploid assembly of any seed-free land plant.

We now have two high-quality *Azolla* genomes: *A. caroliniana* (this study) and *A. filiculoides* (Li et al, 2018). Given the amount of taxonomic uncertainty in this genus, these genomic resources will be invaluable in species delimitation. Surprisingly, our reference genome turned out to be *A. caroliniana* instead of the expected taxon in California, *A. microphylla*, representing either a major expansion of its range (*A. caroliniana* is considered to be restricted to Eastern North America; Flora of North America, 1993) or indicating the necessity for a major taxonomic revision.

We can now tease apart the genomic differences within the two major *Azolla* sect. *Azolla* clades, one of which includes *A. rubra* and *A. filiculoides* and the other that includes *A. microphylla, A. caroliniana*, and *A. mexicana* (Metzgar et al. 2007). In particular, these resources will aid in discerning whether there are any differences between *A. microphylla, A. caroliniana*, and *A. mexicana*, all of which have been reported for California and have at times been treated as synonyms. Knowing what species there are in California will be important for conservation planning as it allows us to sort out the potential invasives that need monitoring and control from the rare natives that warrant conservation attention and hitherto have been unidentifiable. We can now begin to establish barcoding protocols, once the taxonomic situation is resolved, to provide identification resources.

These genomic resources will also provide a foundation for the integration of morphological and microbiome datasets to produce a broad multi-level picture of *Azolla* diversity within California and beyond (Song et al., 2024). Additionally, this chromosome-level genome will be a welcome contribution to fern comparative genomics, which is in its infancy, as only a few chromosome-level assemblies of ferns have currently been published: *Marsilea vestita* (Rahmatpour et al., 2023), *Ceratopteris richardii* (Marchant et al., 2022), *Alsophila spinulosa* (Huang et al., 2022), and *Adiantum capillus-veneris* (Fang et al., 2022). As the only fern included in the CCGP, these genomic resources will provide helpful comparisons to the insights gleaned from the landscape genomics studies of their sister taxon, the seed plants (e.g., Anghel et al. 2022; Mead et al. 2024; Fitz-Gibbon et al. 2023; McEvoy et al. 2023; Huang et al. 2022).

## Acknowledgements

PacBio Sequel II/IIe library prep and sequencing was carried out at the DNA Technologies and Expression Analysis Core at the UC Davis Genome Center, supported by NIH Shared Instrumentation Grant 1S10OD010786-01. Deep sequencing of Omni-C libraries used the Novaseq S4 sequencing platforms at the Vincent J. Coates Genomics Sequencing Laboratory at UC Berkeley, supported by NIH S10 OD018174 Instrumentation Grant. We thank the staff at the UC Davis DNA Technologies and Expression Analysis Core and the UC Santa Cruz Paleogenomics Laboratory for their diligence and dedication to generating high quality sequence data.

## Data Availability

Data generated for this study are available under NCBI BioProject PRJNA720569. Raw sequencing data for sample PRJNA777144 (NCBI BioSample SAMN41564273) are deposited in the NCBI Short Read Archive (SRA). Assembly scripts and other data for the analyses presented can be found at the following GitHub repository: www.github.com/ccgproject/ccgp_assembly

## List of Supplementary Material

**Supp. Fig. 1.**
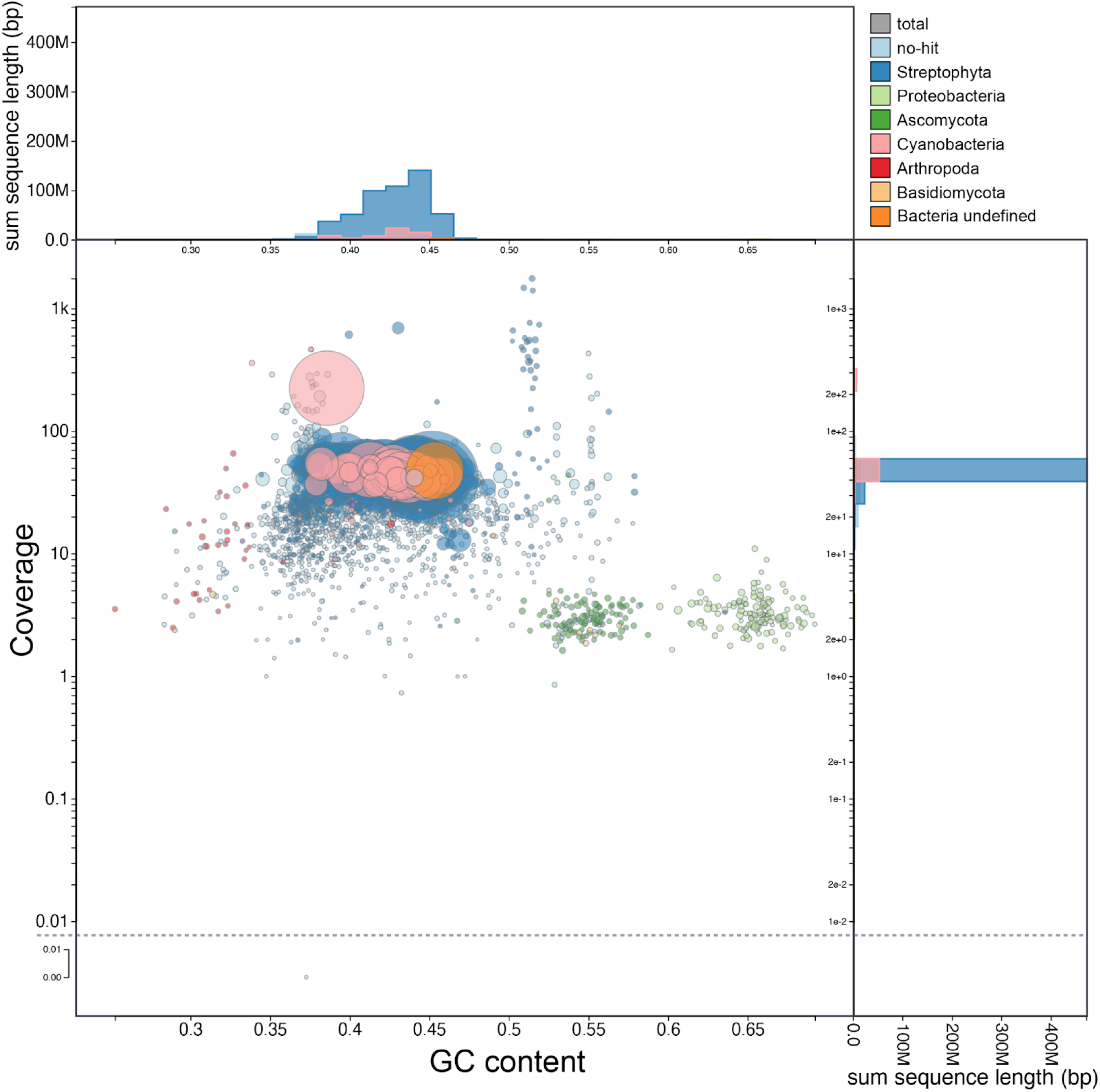
The initial HiFi assembly of *Azolla caroliniana* contained contigs from non-plant organisms. The largest category is from cyanobacteria, represented by the obligate endosymbiont *Nostoc azollae*.

**Supp. Fig. 2.**
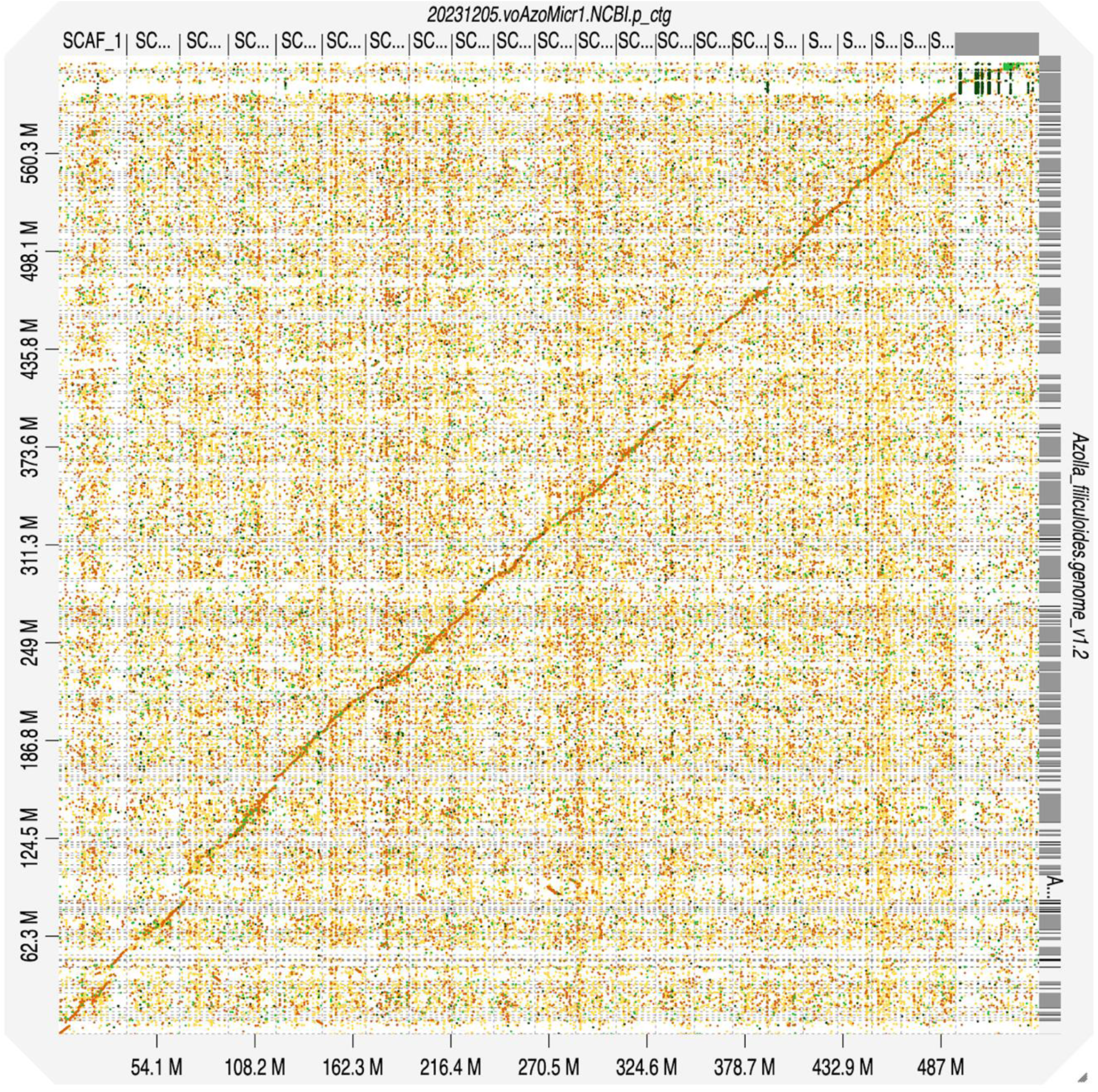
The *Azolla caroliniana* genome (top) is largely collinear with the previously published *A. filiculoides* assembly (side).

**Supp. Fig. 3.**
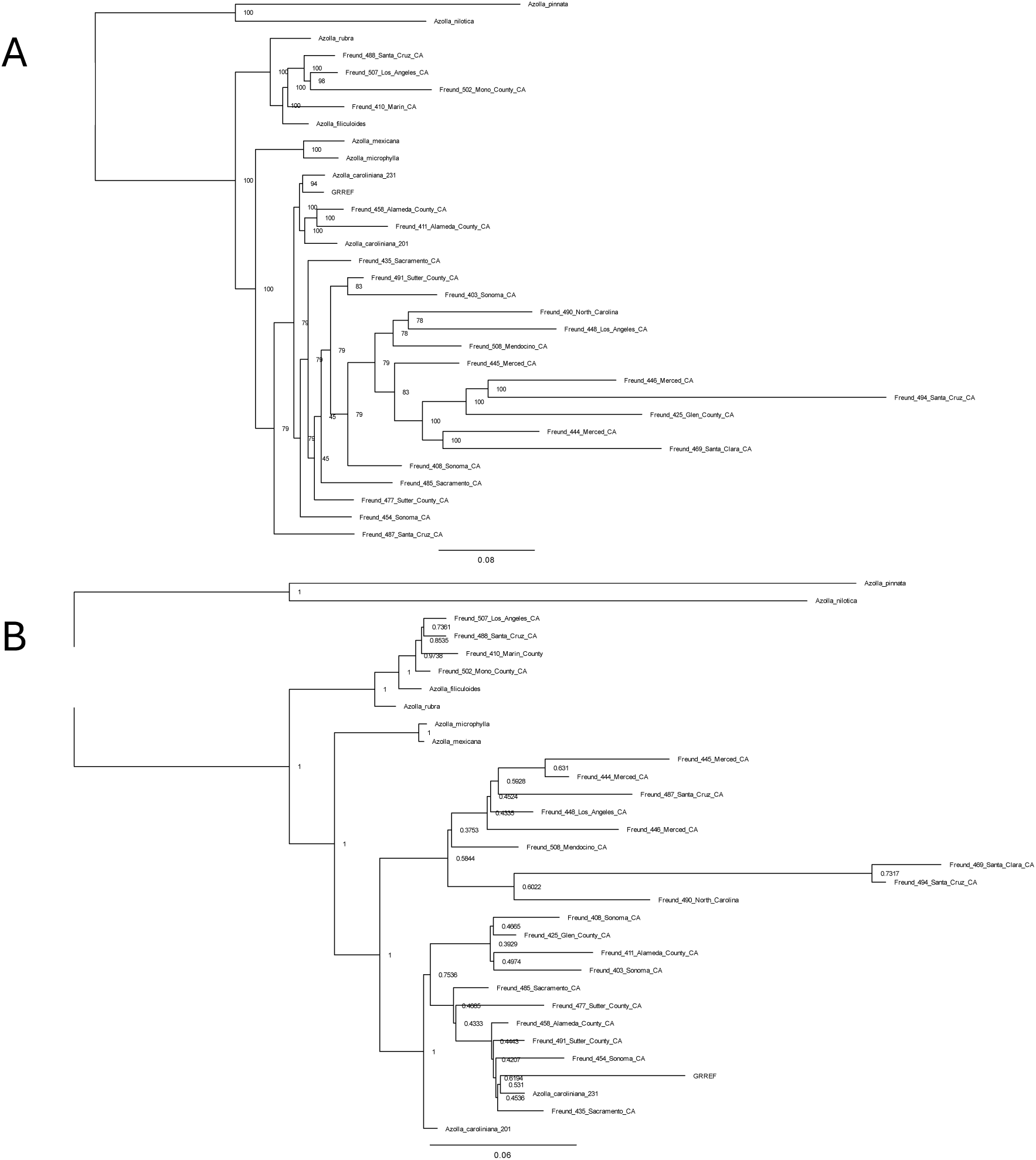
The concatenated-date phylogeny (A) and species tree (B). The branch lengths of the concatenated-date phylogeny maximum likelihood tree are in units of the expected number of substitutions per site; nodes are labeled with ultrafast bootstrap approximation support values from 1000 bootstrap replicates. The branch lengths of the ASTRAL species tree are in coalescent units and nodes are labeled by the support values of the local posterior probability. ASTRAL only estimates branch lengths for internal branches. The *A. caroliniana* reference genome published in this paper is labeled as “GRREF.”

Supplementary Tables 1–3. Detailed taxonomic assignment of removed contigs.

## References

Abdennur, N., and L. A. Mirny. 2020. Cooler: scalable storage for Hi-C data and other genomically labeled arrays. Bioinformatics 36: 311–316.

Anghel, I. G., S. J. Jacobs, M. Escalona, M. P. A. Marimuthu, C. W. Fairbairn, E. Beraut, O. Nguyen, et al. 2022. Reference genome of the color polymorphic desert annual plant sandblossoms, *Linanthus parryae*. Journal of Heredity 113: 712–721.

Beninde, J., E. Toffelmier, and H. B. Shaffer. 2022. A brief history of population genetic research in California and an evaluation of its utility for conservation decision-making W. Murphy. Journal of Heredity 113: 604–614.

Brouwer, P., A. Bräutigam, C. Külahoglu, A. O. E. Tazelaar, S. Kurz, K. G. J. Nierop, A. Van Der Werf, et al. 2014. *Azolla* domestication towards a biobased economy? New Phytologist 202: 1069–1082.

Cabanettes, F., and C. Klopp. 2018. D-GENIES: dot plot large genomes in an interactive, efficient and simple way. PeerJ 6: e4958.

Camacho, C., G. Coulouris, V. Avagyan, N. Ma, J. Papadopoulos, K. Bealer, and T. L. Madden. 2009. BLAST+: architecture and applications. BMC Bioinformatics 10: 421.

Challis, R., S. Kumar, C. Sotero-Caio, M. Brown, and M. Blaxter. 2023. Genomes on a Tree (GoaT): A versatile, scalable search engine for genomic and sequencing project metadata across the eukaryotic tree of life. Wellcome Open Research 8: 24.

Challis, R., E. Richards, J. Rajan, G. Cochrane, and M. Blaxter. 2020. BlobToolKit – Interactive Quality Assessment of Genome Assemblies. G3 Genes|Genomes|Genetics 10: 1361–1374.

Cheng, H., G. T. Concepcion, X. Feng, H. Zhang, and H. Li. 2021. Haplotype-resolved de novo assembly using phased assembly graphs with hifiasm. Nature Methods 18: 170–175.

Cheng, H., Jarvis, E.D., Fedrigo, O., Koepfli, K.P., Urban, L., Gemmell, N.J., Li, H. (2022) Haplotype-resolved assembly of diploid genomes without parental data. Nature Biotechnology, 40:1332–1335. 10.1038/s41587-022-01261-x

Chernomor, O., A. Von Haeseler, and B. Q. Minh. 2016. Terrace Aware Data Structure for Phylogenomic Inference from Supermatrices. Systematic Biology 65: 997–1008.

Danecek, P., J. K. Bonfield, J. Liddle, J. Marshall, V. Ohan, M. O. Pollard, A. Whitwham, et al. 2021. Twelve years of SAMtools and BCFtools. GigaScience 10: giab008.

De Vries, S., and J. De Vries. 2022. Evolutionary genomic insights into cyanobacterial symbioses in plants. Quantitative Plant Biology 3: e16.

Dijkhuizen, L. W., P. Brouwer, H. Bolhuis, G. Reichart, N. Koppers, B. Huettel, A. M. Bolger, et al. 2018. Is there foul play in the leaf pocket? The metagenome of floating fern *Azolla* reveals endophytes that do not fix N _2_ but may denitrify. New Phytologist 217: 453–466.

Evrard, C., and C. Van Hove. 2004. Taxonomy of the American Azolla Species (Azollaceae): A Critical Review. Systematics and Geography of Plants 74: 301–318.

Fang, Y., X. Qin, Q. Liao, R. Du, X. Luo, Q. Zhou, Z. Li, et al. 2022. The genome of homosporous maidenhair fern sheds light on the euphyllophyte evolution and defenses. Nature Plants 8: 1024–1037.

Fiedler, P. L., B. Erickson, M. Esgro, M. Gold, J. M. Hull, J. M. Norris, B. Shapiro, et al. 2022. Seizing the moment: The opportunity and relevance of the California Conservation Genomics Project to state and federal conservation policy. Journal of Heredity 113: 589–596.

Fitz-Gibbon, S., A. Mead, S. O’Donnell, Z.-Z. Li, M. Escalona, E. Beraut, S. Sacco, et al. 2023. Reference genome of California walnut, *Juglans californica*, and resemblance with other genomes in the order Fagales. Journal of Heredity 114: 570–579.

Flora of North America. 2: Pteridophytes and Gymnosperms. 1993. Oxford Univ. Press, New York.

Ghurye, J., M. Pop, S. Koren, D. Bickhart, and C.S. Chin. 2017. Scaffolding of long read assemblies using long range contact information. BMC Genomics 18: 527.

Ghurye, J., A. Rhie, B. P. Walenz, A. Schmitt, S. Selvaraj, M. Pop, A. M. Phillippy, and S. Koren. 2019. Integrating Hi-C links with assembly graphs for chromosome-scale assembly. PLOS Computational Biology 15: e1007273.

Gurevich, A., V. Saveliev, N. Vyahhi, and G. Tesler. 2013. QUAST: quality assessment tool for genome assemblies. Bioinformatics 29: 1072–1075.

Huang, X., W. Wang, T. Gong, D. Wickell, L.-Y. Kuo, X. Zhang, J. Wen, et al. 2022a. The flying spider-monkey tree fern genome provides insights into fern evolution and arborescence. Nature Plants 8: 500–512.

Huang, Y., M. Escalona, G. Morrison, M. P. A. Marimuthu, O. Nguyen, E. Toffelmier, H. B. Shaffer, and A. Litt. 2022b. Reference Genome Assembly of the Big Berry Manzanita (*Arctostaphylos glauca*). Journal of Heredity 113: 188–196.

Inglis, P. W., M. D. C. R. Pappas, L. V. Resende, and D. Grattapaglia. 2018. Fast and inexpensive protocols for consistent extraction of high quality DNA and RNA from challenging plant and fungal samples for high-throughput SNP genotyping and sequencing applications. PLOS ONE 13: e0206085.

Jepson Flora Project. 2024. Jepson eFlora, https://ucjeps.berkeley.edu/eflora/ [accessed on Oct 16, 2024]

Johnson, M. G., L. Pokorny, S. Dodsworth, L. R. Botigué, R. S. Cowan, A. Devault, W. L. Eiserhardt, et al. 2019. A Universal Probe Set for Targeted Sequencing of 353 Nuclear Genes from Any Flowering Plant Designed Using k-Medoids Clustering. Systematic Biology 68: 594–606.

Katoh, K. 2005. MAFFT version 5: improvement in accuracy of multiple sequence alignment. Nucleic Acids Research 33: 511–518.

Kerpedjiev, P., N. Abdennur, F. Lekschas, C. McCallum, K. Dinkla, H. Strobelt, J. M. Luber, et al. 2018. HiGlass: web-based visual exploration and analysis of genome interaction maps. Genome Biology 19: 125.

Korlach, J., G. Gedman, S. B. Kingan, C.-S. Chin, J. T. Howard, J.-N. Audet, L. Cantin, and E. D. Jarvis. 2017. De novo PacBio long-read and phased avian genome assemblies correct and add to reference genes generated with intermediate and short reads. GigaScience 6.

Lee, S., C. R. Bakker, C. Vitzthum, B. H. Alver, and P. J. Park. 2022. Pairs and Pairix: a file format and a tool for efficient storage and retrieval for Hi-C read pairs. Bioinformatics 38: 1729–1731.

Li, F.-W., P. Brouwer, L. Carretero-Paulet, S. Cheng, J. De Vries, P.-M. Delaux, A. Eily, et al. 2018. Fern genomes elucidate land plant evolution and cyanobacterial symbioses. Nature Plants 4: 460–472.

Li, H. 2013. Aligning sequence reads, clone sequences and assembly contigs with BWA-MEM.

Li, H. 2018. Minimap2: pairwise alignment for nucleotide sequences. Bioinformatics 34: 3094–3100.

Li, H., B. Handsaker, A. Wysoker, T. Fennell, J. Ruan, N. Homer, G. Marth, et al. 2009. The Sequence Alignment/Map format and SAMtools. Bioinformatics 25: 2078–2079.

Lumpkin, T. A., and D. L. Plucknett. 1982. Azolla as a green manure: use and management in crop production. Westview Press, Boulder, Colorado.

Madeira, P. T., T. D. Center, J. A. Coetzee, R. W. Pemberton, M. F. Purcell, and M. P. Hill. 2013. Identity and origins of introduced and native Azolla species in Florida. Aquatic Botany 111: 9–15.

Manni, M., M. R. Berkeley, M. Seppey, F. A. Simão, and E. M. Zdobnov. 2021. BUSCO Update: Novel and Streamlined Workflows along with Broader and Deeper Phylogenetic Coverage for Scoring of Eukaryotic, Prokaryotic, and Viral Genomes. Molecular Biology and Evolution 38: 4647–4654.

Marchant, D. B., G. Chen, S. Cai, F. Chen, P. Schafran, J. Jenkins, S. Shu, et al. 2022. Dynamic genome evolution in a model fern. Nature Plants 8: 1038–1051.

McEvoy, S. L., N. Lustenhouwer, M. K. Melen, O. Nguyen, M. P. A. Marimuthu, N. Chumchim, E. Beraut, et al. 2023. Chromosome-level reference genome of stinkwort, *Dittrichia graveolens* (L.) Greuter: A resource for studies on invasion, range expansion, and evolutionary adaptation under global change. Journal of Heredity 114: 561–569.

Mead, A., S. T. Fitz-Gibbon, M. Escalona, E. Beraut, S. Sacco, M. P. A. Marimuthu, O. Nguyen, and V. L. Sork. 2024. The genome assembly of Island Oak (*Quercus tomentella*), a relictual island tree species. Journal of Heredity 115: 221–229.

Metzgar, J. S., H. Schneider, and K. M. Pryer. 2007. Phylogeny and Divergence Time Estimates for the Fern Genus *Azolla*(Salviniaceae). International Journal of Plant Sciences 168: 1045–1053.

Minh, B. Q., H. A. Schmidt, O. Chernomor, D. Schrempf, M. D. Woodhams, A. Von Haeseler, and R. Lanfear. 2020. IQ-TREE 2: New Models and Efficient Methods for Phylogenetic Inference in the Genomic Era. Molecular Biology and Evolution 37: 1530–1534.

Open2C, Abdennur N, Fudenberg G, Flyamer IM, Galitsyna AA, Goloborodko A, Imakaev M, Venev SV. Pairtools: from sequencing data to chromosome contacts. PLOS Computational Biology. 2024 May 29;20(5):e1012164.

Ortiz, E. M., A. Höwener, G. Shigita, M. Raza, O. Maurin, A. Zuntini, F. Forest, et al. 2023. A novel phylogenomics pipeline reveals complex pattern of reticulate evolution in Cucurbitales. bioRxiv: 10.1101/2023.10.27.564367

Rahmatpour, N., L. Kuo, J. Kang, E. Herman, L. Lei, M. Li, S. Srinivasan, et al. 2023. Analyses of *Marsilea vestita* genome and transcriptomes do not support widespread intron retention during spermatogenesis. New Phytologist 237: 1490–1494.

Ramírez, F., V. Bhardwaj, L. Arrigoni, K. C. Lam, B. A. Grüning, J. Villaveces, B. Habermann, et al. 2018. High-resolution TADs reveal DNA sequences underlying genome organization in flies. Nature Communications 9: 189.

Ranallo-Benavidez, T. R., K. S. Jaron, and M. C. Schatz. 2020. GenomeScope 2.0 and Smudgeplot for reference-free profiling of polyploid genomes. Nature Communications 11: 1432.

Rhie, A., S. A. McCarthy, O. Fedrigo, J. Damas, G. Formenti, S. Koren, M. Uliano-Silva, et al. 2021. Towards complete and error-free genome assemblies of all vertebrate species. Nature 592: 737–746.

Rhie, A., B. P. Walenz, S. Koren, and A. M. Phillippy. 2020. Merqury: reference-free quality, completeness, and phasing assessment for genome assemblies. Genome Biology 21: 245.

Shaffer, H. B., E. Toffelmier, R. B. Corbett-Detig, M. Escalona, B. Erickson, P. Fiedler, M. Gold, et al. 2022. Landscape Genomics to Enable Conservation Actions: The California Conservation Genomics Project. Journal of Heredity 113: 577–588.

Sim, S. B., R. L. Corpuz, T. J. Simmonds, and S. M. Geib. 2022. HiFiAdapterFilt, a memory efficient read processing pipeline, prevents occurrence of adapter sequence in PacBio HiFi reads and their negative impacts on genome assembly. BMC Genomics 23: 157.

Simão, F. A., R. M. Waterhouse, P. Ioannidis, E. V. Kriventseva, and E. M. Zdobnov. 2015. BUSCO: assessing genome assembly and annotation completeness with single-copy orthologs. Bioinformatics 31: 3210–3212.

Song, M. J., M. Huynh, S. Lahmeyer, and M. Sedaghatpour. 2023. First Record of the invasive Azolla pinnata subsp. pinnata (Salviniaceae) in California. American Fern Journal 113: 56–57.

Song, M. J., F.-W. Li, F. Freund, C. M. Tribble, E. Toffelmier, C. Miller, H. B. Shaffer, and C. J. Rothfels. 2024. The nitrogen-fixing fern *Azolla* has a complex microbiome characterized by multiple modes of transmission. bioRxiv: doi10.1101/2024.05.20.592813.

Speelman, E. N., M. M. L. Van Kempen, J. Barke, H. Brinkhuis, G. J. Reichart, A. J. P. Smolders, J. G. M. Roelofs, et al. 2009. The Eocene Arctic *Azolla* bloom: environmental conditions, productivity and carbon drawdown. Geobiology 7: 155–170.

Stergianou, K. K., and K. Fowler. 1990. Chromosome numbers and taxonomic implications in the fern genusAzolla (Azollaceae). Plant Systematics and Evolution 173: 223–239.

Tillich, M., P. Lehwark, T. Pellizzer, E. S. Ulbricht-Jones, A. Fischer, R. Bock, and S. Greiner. 2017. GeSeq – versatile and accurate annotation of organelle genomes. Nucleic Acids Research 45: W6–W11.

Toffelmier, E., J. Beninde, and H. B. Shaffer. 2022. The phylogeny of California, and how it informs setting multispecies conservation priorities. Journal of Heredity 113: 597– 603.

US Federal Noxious Weed List. United States Department of Agriculture.

Watanabe, I., Ma. T. Lapis-Tenorio, T. S. Ventura, and B. C. Padre. 1993. Sexual hybrids of *Azolla filiculoides* with *A. microphylla*. Soil Science and Plant Nutrition 39: 669– 676.

Wolff, J., R. Backofen, and B. Grüning. 2022. Loop detection using Hi-C data with HiCExplorer. GigaScience 11: giac061.

Workman, R., R. Fedak, D. Kilburn, S. Hao, K. Liu, and W. Timp. 2018. High Molecular Weight DNA Extraction from Recalcitrant Plant Species for Third Generation Sequencing v1.

Zhang, C., M. Rabiee, E. Sayyari, and S. Mirarab. 2018. ASTRAL-III: polynomial time species tree reconstruction from partially resolved gene trees. BMC Bioinformatics 19: 153.

